# *HvbZIP33* and *HvbZIP76* have overlapping roles in foliar transpiration in drought-stressed barley

**DOI:** 10.64898/2026.01.02.697348

**Authors:** Asis Shrestha, Michael Schneider, Götz Hensel, Corinna Sauer, Jan Benndorf, Thuy Huu Nguyen, Suresh Venkata Bonthala, Jochen Kumlehn, Benjamin Stich, Jens Léon, Ali Ahmad Naz

## Abstract

Basic leucine zippers (bZIP) constitute one of the biggest protein families and evolutionarily conserved transcription factors (TFs) in plants. We obtained mutant lines for two bZIP TFs, *HvbZIP33* and *HvbZIP76*, in the genetic background of the barley cultivar Golden Promise (GP) via targeted gene-specific mutagenesis using an RNA-guided Cas9 endonuclease. A comprehensive morphological, physiological and transcriptomic analysis was performed in wild-type GP compared with *hvbzip33* and *hvbzip76* mutants under drought stress. The morphological and physiological changes were similar in both mutants and in the wild-type GP. Most strikingly, the mutants exhibited accelerated wilting and increased water loss. This effect was primarily caused by higher stomatal conductance (*g_s_*) and transpiration rate (*E*) in mutants compared to wild-type GP under both control and drought conditions, which in turn had a detrimental effect on the mutants’ intrinsic water use efficiency (*iWUE*). Likewise, the transcriptome profiles of *hvbzip33* and *hvbzip76* were more similar to each other than those of wild-type GP. We found that the number of differentially regulated genes under control versus drought-stress conditions was higher in the mutants than in wild-type GP, suggesting that the mutants try to compensate for accelerated foliar water loss. The study highlights the essential roles of *HvbZIP33* and *HvbZIP76* in balancing water loss in barley. These findings provide a foundation for engineering enhanced drought tolerance in barley through targeted manipulation of these genes to optimize transpiration rates.

## 1. Introduction

Drought stress is one of the most significant challenges of food production in the current context of climate uncertainty (Leng and Hall, 2019). Hence, drought stress tolerance is also one of the widely researched topics in plant science and it is now well established that tolerance mechanisms involve a complex interaction of multiple gene regulatory processes (Nakashima *et al*., 2025). Transcription factors (TFs) are central to these gene expression networks that connect drought signaling to stress-responsive genes to determine optimal physiological, biochemical and growth adjustments (Dhatterwal *et al*., 2024). Among them, the basic leucine zipper (bZIP) family is one of the most prominent and evolutionarily conserved TF families in plants. bZIP TFs have two conserved domains, namely a basic region and a leucine zipper domain (Jakoby *et al*., 2002). The basic region is highly conserved and facilitates the binding of bZIP proteins to specific DNA sequences, typically containing a core ACGT motif, such as G-box (C**ACGT**G), C-box (C**ACGT**C) and A-box (T**ACGT**A) sequences (Li *et al*., 2023). Beyond these ACGT-core motifs, the potential non-ACGT targets of bZIP proteins are also identified in cereals and are referred to as coupling element 1 (CE1, CCACCG) (Shen and Ho, 1995; Shen *et al*., 1996; Roychoudhury and Sengupta, 2009) and CE3 (CGTGTC) (Hobo *et al*., 1999). The leucine zipper region is another conserved binding domain that is characterized by leucine residues, which facilitate the formation of homo- or heterodimers with different members of the bZIP family proteins (Kaur *et al*., 2025). This modular architecture of bZIP TFs highlights their versatility in regulating a wide range of biological processes in plants.

Ten groups of bZIP proteins (A, B, C, D, E, F, G, H, I and S) have been described for Arabidopsis and members of the group A are involved in responses to abiotic stresses, including drought, cold and salinity tolerance (Fujita *et al*., 2005; Yoshida *et al*., 2015; Chang *et al*., 2019). They are usually referred to as ABA-responsive element (ABRE)-binding TFs (ABFs). Among them, the binding property of four ABFs (ABF1, AREB1/ ABF2, ABF3 and AREB2/ ABF4) to ABREs was identified through yeast one-hybrid screening and electrophoretic mobility shift assays (Choi *et al*., 2000; Uno *et al*., 2000). These four respective genes are highly expressed in vegetative tissue and their expression is significantly induced by abiotic stress (Yoshida *et al*., 2010; Yoshida *et al*., 2015). Over-expression lines of ABFs showed enhanced stress adaptation while the knockout lines were sensitive to drought stress in several plant species, including Arabidopsis (Fujita *et al*., 2005; Yoshida *et al*., 2010; Hossain *et al*., 2010; Huang *et al*., 2010; Amir Hossain *et al*., 2010; Yoshida *et al*., 2015; Wang *et al*., 2016). Four other genes from the AREB family (*ABI5*/*DPBF1*, *DPBF2*, *AREB3*/*DPBF3* and *DPBF4*) have more specialized expression patterns. Among the four *DPBF* genes, *DPBF1* is expressed in roots, leaves and reproductive tissues, while *DPBF2* expression is strictly specific to seeds. *DPBF3* shows lower overall expression across tissues and growth conditions than other *DPBF* genes. Unlike stress-inducible *ABFs*, the four *DPBFs* are involved in seed germination, seed maturation, floral development and the regulation of energy homeostasis (Nijhawan *et al*., 2008; Alonso *et al*., 2009; van Leene *et al*., 2016).

Despite the importance of group A bZIP TFs in stress adaptation and plant development, a comprehensive understanding of their functional diversity and the regulatory networks that govern their actions remains incomplete. This is especially valid for barley and concerns both the downstream target genes they activate and the upstream signals that modulate their expression and activity. For instance, the roles of some genes of the ABF and DPBF class of bZIP TFs were studied in rice, wheat and maize (Xiang *et al*., 2008; Hossain *et al*., 2010; Tang *et al*., 2012; Wei *et al*., 2012; Liu *et al*., 2014; Zhang *et al*., 2017; He *et al*., 2024). For example, multiple bZIP TFs in rice, namely, *OsABF2*, *OsbZIP23*, *OsbZIP46*, *OsbZIP71* and *OsbZIP72,* act as positive regulators of drought and salt tolerance by integrating ABA signaling. Likewise, bZIP genes in wheat (sub-group A) conferred abiotic stress tolerance when over-expressed in Arabidopsis (*Traes_7AL_25850F96F.1*, *TabZIP174*), as well as nitrogen use efficiency (*TabZIP60*) in wheat (Li *et al*., 2016; Yang *et al*., 2019; Agarwal *et al*., 2019). To date, only one group A bZIP gene has been functionally characterized in barley. Recent studies showed that constitutive overexpression of *HvbZIP77* in Arabidopsis enhanced drought tolerance but reduced seed size (Al-Sayaydeh *et al*., 2024).

From the genome-wide identification of bZIP genes in barley, it was concluded that they belong to a large family with 89 and 92 genes, as reported for the Morex genome versions 1 (Pourabed *et al*., 2015) and 2 (Zhong *et al*., 2021), respectively. These members of the bZIP family were grouped into ten clusters (A-I and S) based on the gene structure and expression pattern. Although these studies provided a comprehensive catalog and classification of bZIP genes in barley, their biological roles remain largely unexplored. Previous evidence from promoter activation assays in barley and other cereals suggests that group A bZIP transcription factors (*HvABI5/HvbZIP56, HvABF1/ HvbZIP33, HvABF2/ HvbZIP77* and *HvABF3/ HvbZIP52*) may act as positive regulators of ABA-responsive genes. For example, minimal promoter regions of *HVA22* containing an ABA-responsive complex (ABRC) composed of ACGT-ABRE, CE3 and CE1 have been shown to drive ABA-dependent reporter expression in barley (Shen *et al*., 1993; Shen and Ho, 1995; Gómez-Cadenas *et al*., 1999; Casaretto and Ho, 2003; Shen *et al*., 2004; Cao *et al*., 2007). Later, Schoonheim *et al*. (2009) showed that ectopic expression in alurone cells co-transfected with *ABF1/ABF3* and ABRC:GUS was sufficient to transactivate GUS even without ABA application.

Nonetheless, the functional significance of most bZIP TFs in barley remains unknown. This research aimed to identify and characterize bZIP TFs in barley using a combination of bioinformatics approaches and experimental validation. We employed Cas9-mediated genome editing to create potential loss-of-function mutants of two bZIP TFs in barley. Subsequently, the mutant lines were phenotyped under drought-stress and control conditions at the seedling stage. A comprehensive RNA-seq analysis was performed to profile the transcriptomes of mutant lines for two bZIP TFs under drought stress to identify their downstream target genes.

## 2. Materials and Methods

### 2.1. Selection of bZIP transcription factors for functional studies

In the first step, we compared the bZIP genes annotated in the barley reference genome (Morex V3) with those present in Morex V2 (Zhong *et al*., 2021). Because the goal was to identify barley homologs of AREBs (class A bZIP family proteins) from Arabidopsis, we aligned the protein sequence of the HvbZIPs in the Morex V3 sequence and nine AREBs (class A bZIP family proteins) from Arabidopsis using the “msa” package in R. The protein alignment file was used as input to construct the phylogenetic tree using the “phytools” package in R.

Among the genes that clustered together with Arabidopsis AREBs, we selected HORVU.MOREX.r3.6HG0619650, *HvbZIP76* and HORVU.MOREX.r3.3HG0300770, *HvbZIP33* for functional validation based on prior information on those genes (Figure S1). We selected a target motif in exon 1 of *HvbZIP33* and *HvbZIP76* (Figure 1). The respective gRNA was synthesized and inserted into pSH91 using a restriction enzyme (*BsaI*). The validated clones were digested with *SfiI* and the gRNA and Cas9-containing fragment was transferred to the generic binary vector p6ID35STE9 (DNA Cloning Service, Hamburg), which carries *hpt* for plant selection. Immature embryos of barley inbred Golden Promise (GP) were used for *Agrobacterium*-mediated DNA transfer according to (Marthe *et al*., 2015).

**Figure 1:**
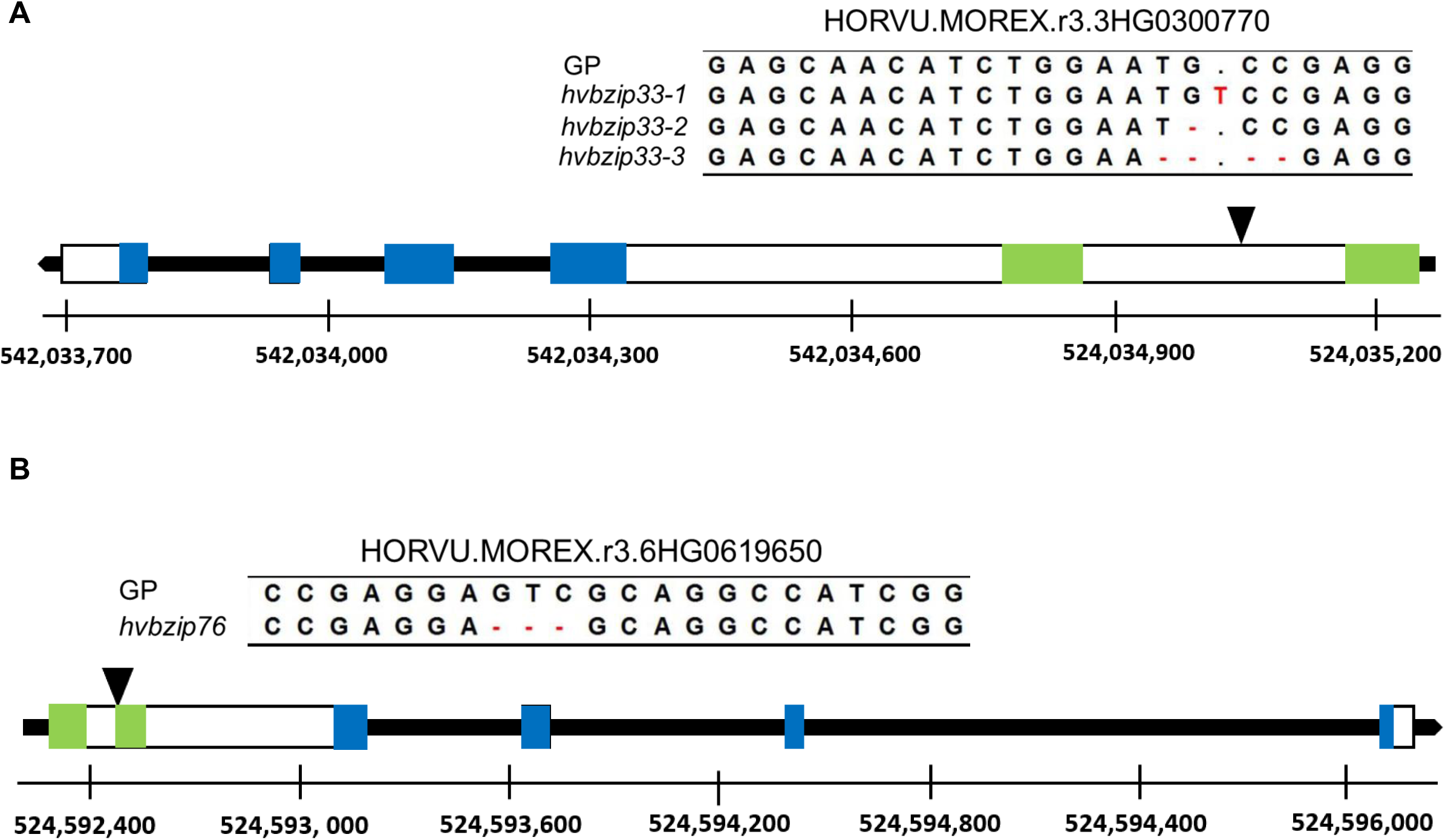
Mutation even in the coding sequence of *HvbZIP33* and *HvbZIP76*. Insertion and deletion mutations were detected in (A) *HvbZIP33* and (B) *HvbZIP76* in genome editing pipelines using the RNA-guided Cas9 endonuclease. The rectangular boxes are the exons; the green and blue regions represent conserved domains. The black triangle indicates the site of mutation.

### 2.2. Genotyping the T_0_, T_1_ and T_2_ generation

Mutation events in T0 progenies were detected by Sanger sequencing at both the target and predicted off-target sites. Subsequently, 48 T_1_ progenies from T_0_ plants that exhibited mutation events were screened for T-DNA. T_1_ progeny from each selected T_0_ line that tested negative for T-DNA insertion were sequenced to detect homozygous mutation events. The positive T_1_ progeny for mutation events were selfed to obtain T_2_ seeds. Finally, T_2_ progenies were again genotyped for the absence of T-DNA as a confirmation and sequenced to identify transgene-free T_2_ plants that are homozygous for mutation events in the target genes. Verified T_2_ progenies were self-pollinated to obtain seeds from individuals homozygous for mutant alleles, enabling characterization of the mutants under drought stress. The mutant lines for *HvbZIP33* and *HvbZIP76* are referred to as *hvbzip33* and *hvbzip76* in the following text.

### 2.3. Plant growth conditions and drought treatment

Seeds were pregerminated using a peat-based potting mixture, ED73 classic, produced and marketed by Einheitserde, Germany and two-day-old seedlings were transferred to a pot (8 x 8 x 7 cm). The pots were filled with an equal volume of mixture containing 60% natural sand and 40% topsoil (Terrasoil; Cordel and Sohn). The plants were grown in the greenhouse under the following conditions: a daily mean temperature of 18-22°C, a 14/10 h light/dark cycle and a light intensity of 110-150 μmol m^-2^ s^-1^. The field capacity of the soil was maintained at 80% under control conditions. Pots were weighed twice a day and manually watered to maintain a constant soil moisture level. Drought stress was induced by withholding water from 14-day-old seedlings. The weight of the pots was recorded daily to ensure a consistent moisture level in all pots.

### 2.4. Tissue water status and oxidative stress marker evaluation

Relative water content (RWC) and electrolyte leakage (EL) were evaluated according to (Shrestha *et al*., 2022). First, the tip of the first fully expanded leaf from the top was removed. Then, two leaf sections (around 2 cm each) were detached from the first fully expanded leaf and the fresh weight was recorded (FW). Subsequently, the leaf sections were dipped into centrifuge tubes containing 10 ml of deionized water and incubated at room temperature for 24 h. The leaf sections were removed from the centrifuge tube and excess water was gently wiped away with a paper towel before measuring the turgor weight (TW). The dry weight was recorded after the sample was oven-dried at 70°C for 72 hours. RWC was estimated as (FW-DW)/(TW-DW) *100.

For EL measurement, centrifuge tubes were filled with 10 mL of deionized water and the initial electrical conductivity (ECi) was recorded. Then, two leaf sections (approximately 2 cm each) from the same leaf used for RWC were excised and placed in a centrifuge tube containing 10 mL of double-distilled water. The tubes were stored in the dark at room temperature. Electrical conductivity was measured after a 24 h rehydration period (ECf). After the final reading, the samples were boiled at 100°C for 30 minutes, cooled to room temperature and the total electrical conductivity (ECt) was measured. EL was expressed as (ECf-ECi)/(ECt-ECi)*100.

Oxidative damage to the lipid membrane during drought was estimated by determining the malondialdehyde (MDA) concentration using the thiobarbituric acid (TBA) method (Hodges *et al*., 1999), adapted to a microplate-based protocol (Dziwornu *et al*., 2018) with some modifications. Shoot samples were homogenized in liquid nitrogen and MDA was extracted with 1.5 ml of 0.1% trichloroacetic acid (TCA), followed by centrifuging at 14,000 *g* for 15 minutes at 4°C. Then, 500 μL of supernatant was mixed with reaction solution I (0.01% 2,6-di-tert-butyl-4-methylphenol (BHT) in 20% TCA) and reaction solution II (0.65% TBA, 0.01% BHT in 20% TCA) in a 1:1 ratio. Reaction and sample mix were incubated at 95°C for 30 minutes. The reaction was stopped on ice for five minutes and the mixture was centrifuged at 8000 *g* for 10 minutes at 4°C. The absorbance was measured at 440 nm, 532 nm and 600 nm using a microplate reader (TECAN Infinite 200 Pro, TECAN Group Limited, Switzerland). MDA concentration was expressed as nanomoles per gram fresh weight.

### 2.5. Gas exchange experiment

Gas exchange was measured using a portable photosynthesis system (LI-6400XT gas exchange analyzer; LI-COR Biosciences) 3, 6 and 9 days after the start of the drought stress treatment (DAS). The measurements were made on the first fully expanded leaves from the top at 3 DAS (three true leaves fully expanded). Subsequently, measurements were made at 6 and 9 DAS on the same leaf for all plants in each treatment to minimize leaf effects. The parameters inside the leaf chamber were set to a saturated photosynthetic active radiation of 1500 μmol m^−2^ s^−1^, a relative humidity of approximately 52%, a temperature corresponding to the leaf temperature and a flow rate of 400 μmol s^−1^. The infrared gas exchange analyzer automatically logs the photosynthetic parameters, including the rate of CO_2_ assimilation (*A*), intercellular CO_2_ (*C_i_*), stomatal conductance (*g_s_*) and transpiration rate (*E*). All observations were adjusted to 25°C. The experiment was performed twice, with four biological replications per genotype per treatment (i.e., eight plants).

### 2.6. Data processing and statistical analysis

Statistical significance was assessed using open-access statistical computing software R, version 4.3.1. The analysis of variance (ANOVA) for physiological and photosynthetic traits was performed using the following model

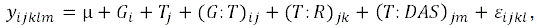

where *y_ijklm_* was the phenotypic observation, (μ) the general mean, (*G_i_*) the effect of genotype, (*T_j_*) the effect of treatment, (*G: T*)_*ij*_ the effect of genotype by treatment interaction, (*T: R*)_*jk*_ the effect of the experiment nested within treatment, (*T: DAS*)_*jl*_ effect of days after the start of stress nested within treatment and (*d_ijkl_*) the error term. Because plant biomass was only estimated at 9 DAS, the ANOVA was performed for this character using the above equation without (*T: DAS*)_*jl*_ effect.

Adjusted entry means and standard errors of means were estimated using the “emmean” package in R. Multiple mean comparisons were performed using the LSD test.

### 2.7. RNAseq analysis of *hvbzip76* and *hvbzip33* under drought stress

We profiled the transcriptomes of GP, *hvbzip33-3 and hvbzip76* under both control and drought stress conditions. The growing environment and stress treatment were identical to those described previously for the physiological experiments. Fresh shoot samples were collected 7 DAS, flash-frozen in liquid nitrogen and stored at −80°C before RNA extraction.

Shoot materials were homogenized in liquid nitrogen, and RNA was extracted using the Monarch RNA Miniprep Kit (New England Biolabs, USA) according to the manufacturer’s instructions. The RNA concentration and quality were determined by electrophoresis on a 1% Agarose gel and by NanoDrop (NanoDrop 2000c, Thermo Fisher Scientific, USA) before shipping. RNA integrity was further verified by using a Bioanalyzer (Agilent 2100, Agilent Technologies, USA). The samples that passed the RNA integrity check (with an integrity value of≥6.3) were used for library preparation. Novogen Europe performed library preparation and sequencing.

We obtained the paired-end read sequence and the quality control of raw sequencing reads was performed using FastQC (Andrews, 2010). Then, the adaptor sequence and reads less than 70 bp were trimmed using Trimmomatic (Bolger *et al*., 2014). Trimmed reads were aligned to the barley reference genome sequence (https://data.jgi.doe.gov/refine-download/phytozome?organism=HvulgareMorex&expanded=Phytozome-702) using HISAT2 (Pertea *et al*., 2016). The unmapped reads were filtered before further processing. To be selected as a mapped read, ≥80% of the length of a read should have mapped to the reference sequence sharing ≥ 90% identity. On average, more than 80% of the trimmed reads mapped to the reference. Differential gene expression (DEG) analysis was performed using the edgeR package (Chen *et al*., 2016). First, the read count for uniquely mapped sequences was normalized to the sample’s sequencing depth and expressed as log counts per million mapped reads. A negative binomial distribution was fitted prior to DEG estimation using the log2 fold change in read counts for the drought treatment relative to the control. Quasi-likelihood F-test was applied and *P* ≤ 0.05 was set as a significance threshold (Chen *et al*., 2016). Multiple testing correction was performed for the significant DEGs list to control the false discovery rate to *P* ≤ 0.05, according to (Benjamini and Hochberg, 1995).

Gene ontology (GO) enrichment analysis (GOEA) for DEGs was performed using goatools v1.4.12 (Klopfenstein *et al*., 2018) to identify the enriched GO terms. For this analysis, we annotated the whole proteome of Morex using InterProScan5 (Jones *et al*., 2014). We used the GO annotation of the entire barley proteome as a background set in GOEA.

## 3. Results

### 3.1. Mutation events in *HvbZIP33* and *HvbZIP76*

Recently, (Zhong *et al*., 2021) showed that the Morex genome (version 2) comprises 92 *HvbZIP* genes. We identified five additional members beyond those listed by (Zhong *et al*., 2021) in the Morex reference version 3 (Table S1). Phylogenetic analysis revealed that 18 HvbZIP TFs belong to group A bZIP proteins, eight more than those reported for Arabidopsis. The 18 genes of group A can be assigned to four sub-clusters: ABF1-ABF4-like sub-cluster, DPBF3-DPBF4-like sub-cluster, DPBF1-DPBF2-like sub-cluster and bZIP15-like sub-cluster (Figure S1). The first sub-cluster of group A bZIPs that branched alongside ABF1-ABF4 comprised four genes: *HvbZIP56*, *HvbZIP76*, *HvbZIP77* and *HvbZIP96*. Four *HvbZIPs*, including *HvbZIP33,* clustered alongside the DPBF1-DPBF2-like sub-cluster. *HvbZIP30* and *HvbZIP07* were the closest barely proteins to the AtDPBF3-AtDPBF4-like sub-cluster, whereas eight *HvbZIPs* were close to Arabidopsis *bZIP15* (Figure S1). The genes used for functional validation in the current study, *HvbZIP33* and *HvbZIP76*, comprise four exons each. The N-terminal region consists of two adjacent intrinsically disordered regions, including phosphorylation sites and binding sites for co-activators and repressors. The C-terminal region contains the conserved bZIP domain that is responsible for dimerization and DNA binding (Figure 1).

After analyzing the regenerated plants, we obtained seven transgene-positive plants carrying gRNAs targeting *HvbZIP33* and one transgene-positive plant carrying a gRNA targeting *HvbZIP76* (Table S2). Sequencing of target regions detected mutation events in six T_0_ progenies for *HvbZIP33* and one for *HvbZIP76* (Table S2). No off-target mutations were detected at predicted off-target sites for both gRNAs. We detected three allelic mutants for *HvbZIP33* with one bp insertion, one bp deletion and four bp deletion, namely *hvbzip33-1*, *hvbzip33-2* and *hvbzip33-3* (Figure 1A). All three allelic mutations introduced premature stop codons upstream of the bZIP domain in the HvbZIP33 protein, thereby completely disrupting the bZIP motif in all mutants (Figure S2). We identified a single allelic variant of *HvbZIP76*, characterized by a GTC deletion in the first exon (Figure 1B). The three bp deletion resulted in an in-frame removal of a serine residue within the intrinsically disordered region (Figure S3). This allelic mutant is referred to as *hvbzip76* in the following text.

### 3.2. Physiological and biochemical response of *hvbzip33* and *hvbzip76* to drought stress

To understand the functions of *HvbZIP33* and *HvbZIP76*, we compared wild-type Golden Promise (GP) and the mutants with respect to morphological, physiological and biochemical traits under control and drought conditions. Plant height and shoot fresh weight at the seedling stage (21-day-old plants) were highest for GP and lowest for *hvbzip33-3* under control conditions (Figure 2A-B). We observed a decrease in plant height and shoot weight in young seedlings in a treatment-dependent manner (Figure 2). Upon exposure to drought stress, all allelic mutants of *hvbzip33* and *hvbzip76* showed wilting symptoms considerably earlier than GP. Initial wilting symptoms appeared at 6 DAS, with pronounced differences between GP and mutants occurring at 7 DAS (Figure 3a). Measurements of RWC and EL revealed significant effects of genotype, treatment and the interaction between genotype and treatment (Table 1). A significant reduction in RWC was observed at 9 DAS, which was more pronounced in *hvbzip33* and *hvbzip76* mutants compared to GP under drought stress (Figure 3b). EL, as an indicator of membrane damage, also increased under drought stress, with EL values significantly greater in *hvbzip33-3* than wild-type GP at 9 DAS (Figure S4a). Although MDA concentration rose under drought stress relative to control conditions at 9 DAS, no significant differences were detected between wild-type GP and any of the *hvbzip76* and *hvbzip33* mutants (Figure S4b). We did not observe differences in plant morphology between adult wild-type GP plants and mutants under control conditions (Figure S5).

**Figure 2:**
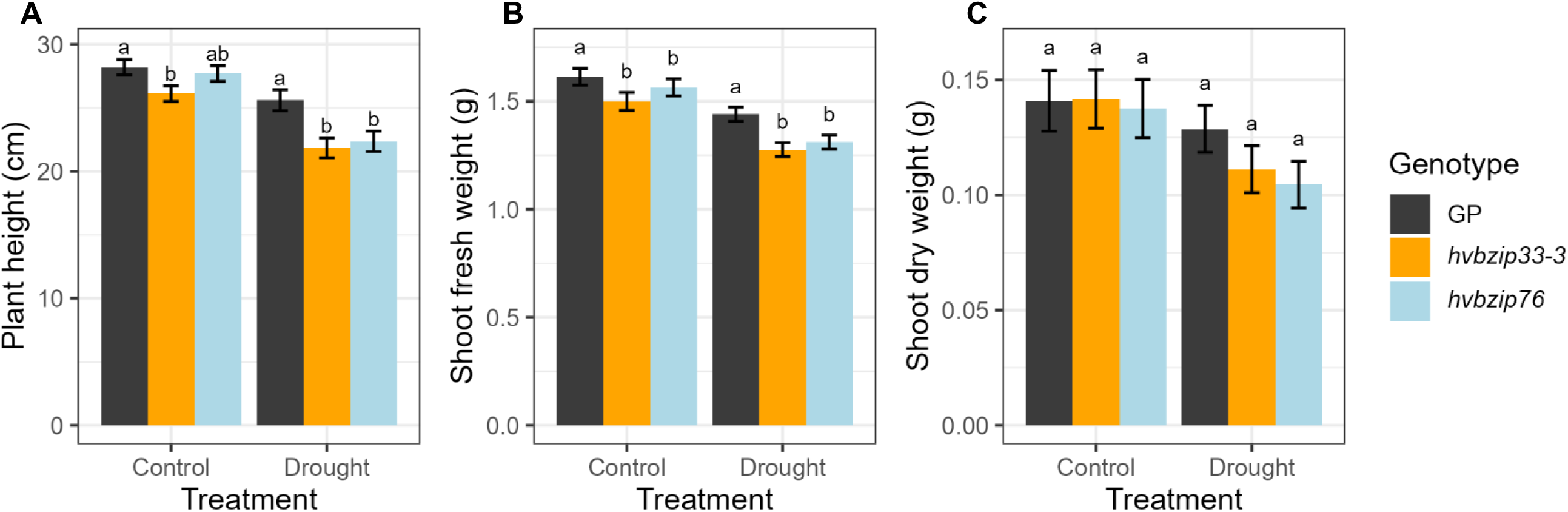
Morphological parameters evaluated under control and drought conditions in Golden Promise (GP) and mutants. The effect of drought stress on (A) plant height, (B) shoot fresh weight, and (C) shoot dry weight. Drought stress was imposed on 14-day-old seedlings via controlled dehydration, ensuring uniform water loss across all experimental units. The morphological evaluation was performed 9 days after the start of stress treatment. Two independent experiments were performed. Each experiment comprised four biological replications per genotype per treatment. The bar represents mean ± standard error (n = 8 to 12). Indexed letter represents significant differences (p ≤ 0.05) between the genotypes using the LSD multiple mean comparison test.

**Figure 3:**
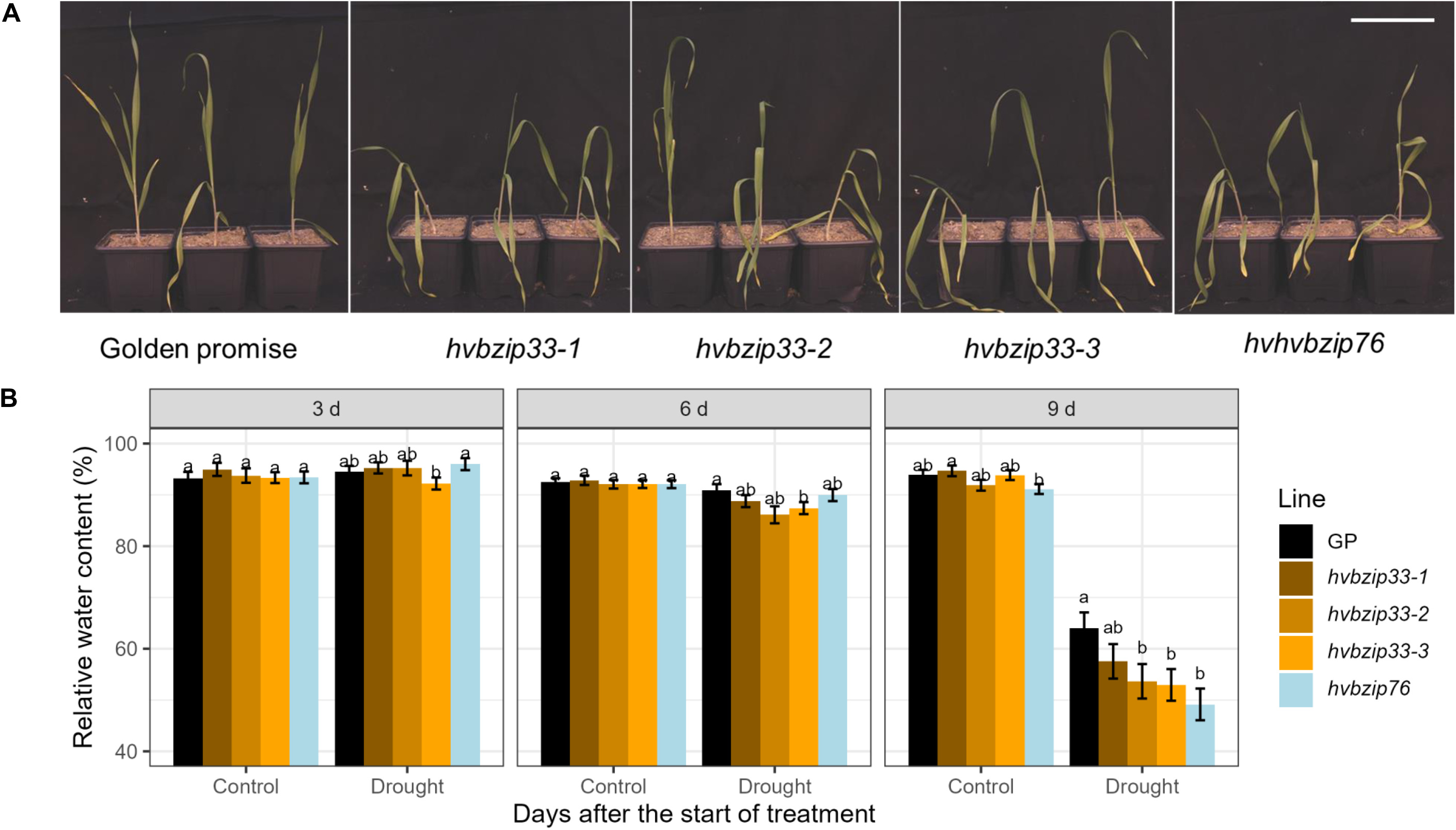
Foliar relative water content evaluated under control and drought conditions in Golden Promise (GP) and mutants. (A) The effect of drought stress on Golden promise (GP) and *hvbzip33* and *hvbzip76* mutants 7 days after stress (DAS). White scale bar corresponds to a length of 8 cm. (B) Relative water content in the first fully expanded leaf from the top at 3, 6 and 9 days after the start of stress treatment (DAS). Drought stress was imposed on 14-day-old seedlings via controlled dehydration, resulting in a uniform water loss across all experimental units. The bar represents mean ± standard error (n = 8 to 12). Indexed letter represents significant differences (p ≤ 0.05) between the genotypes using the LSD multiple mean comparison test.

**Table 1:** Analysis of variance in physiological and morphological traits evaluated under control and drought conditions. Drought stress was imposed on 14-day-old seedlings via controlled dehydration, resulting in a uniform water loss across all experimental units. Photosynthetic and physiological traits were evaluated 3, 6, and 9 days after the start of the stress treatment (DAS). Morphological data were collected at the end of the experiment, 9 days after stress. Asterisks indicate level of significance (***, p ≤ 0.001; **, p ≤ 0.01 *, p ≤ 0.05, not significant (ns), p > 0.05). The experiment was repeated twice (Repetition), with four to six biological replicates per repetition. Abbreviation: not applicable (NA).

| Traits | Genotype | Treatment | Genotype:Treatment | Treatment:Repitition | Treatment:DAS |
| --- | --- | --- | --- | --- | --- |
| $C_i$ | *** | *** | ns | *** | *** |
| $A$ | ns | *** | ns | *** | *** |
| $g_s$ | *** | *** | ns | *** | *** |
| $E$ | *** | *** | ns | ** | *** |
| $iWUE$ | *** | *** | * | ns | ns |
| RWC (%) | ** | *** | * | *** | *** |
| MDA (mg g<sup>-1</sup>) | ns | *** | ns | NA | *** |
| EL (%) | ** | *** | * | *** | *** |
| Plant height (cm) | *** | *** | ns | ns | NA |
| Shoot fresh weight (g) | *** | *** | ns | *** | NA |
| Shoot dry weight (g) | ns | * | ns | ns | NA |

### 3.3. Photosynthetic response of *hvbzip33* and *hvbzip76* to drought stress

Next, we conducted a gas-exchange experiment to quantify differences in photosynthetic parameters among wild-type GP, *hvbzip33-3* and *hvbzip76*. In general, under control conditions, *g_s_*, *E* and *C_i_* were slightly lower in wild-type GP than in both mutant lines (Figure 4A-B, Figure S6). *E* and *g_s_* decreased under drought stress, with the reduction being more pronounced in wild-type GP than in *hvbzip33-3* and *hvbzip76* at 3 and 6 DAS. However, the pattern was reversed at 9 DAS under drought (Figure 4A-B). Without drought stress, *A* was only slightly higher (statistically non-significant) in mutants *hvbzip33-*3 and *hvbzip76* than in wild-type GP. At 3 DAS, *hvbzip76* had shown the highest *A* value under drought stress. Although statistically non-significant, *A* was lower in both mutants than in wild-type GP at 9 DAS under drought stress (Figure 4C). We also estimated *iWUE* as the ratio of *A* to *g_s_*. Because GP transpired less than both mutants, *iWUE* in wild-type GP was significantly higher under control conditions compared to the mutants (Figure 4D). Likewise, *iWUE* increased significantly under drought stress compared to control conditions, in a genotype-dependent manner. For example, we observed a clear trend of lower *iWUE* at all three measurement time points in *hvbzip33-3* than in wild-type GP. In contrast, *iWUE* was lower in *hvbzip76* than in wild-type GP under control conditions, whereas it was on a par in wild-type GP and *hvbzip76* under drought stress (Figure 4D).

**Figure 4:**
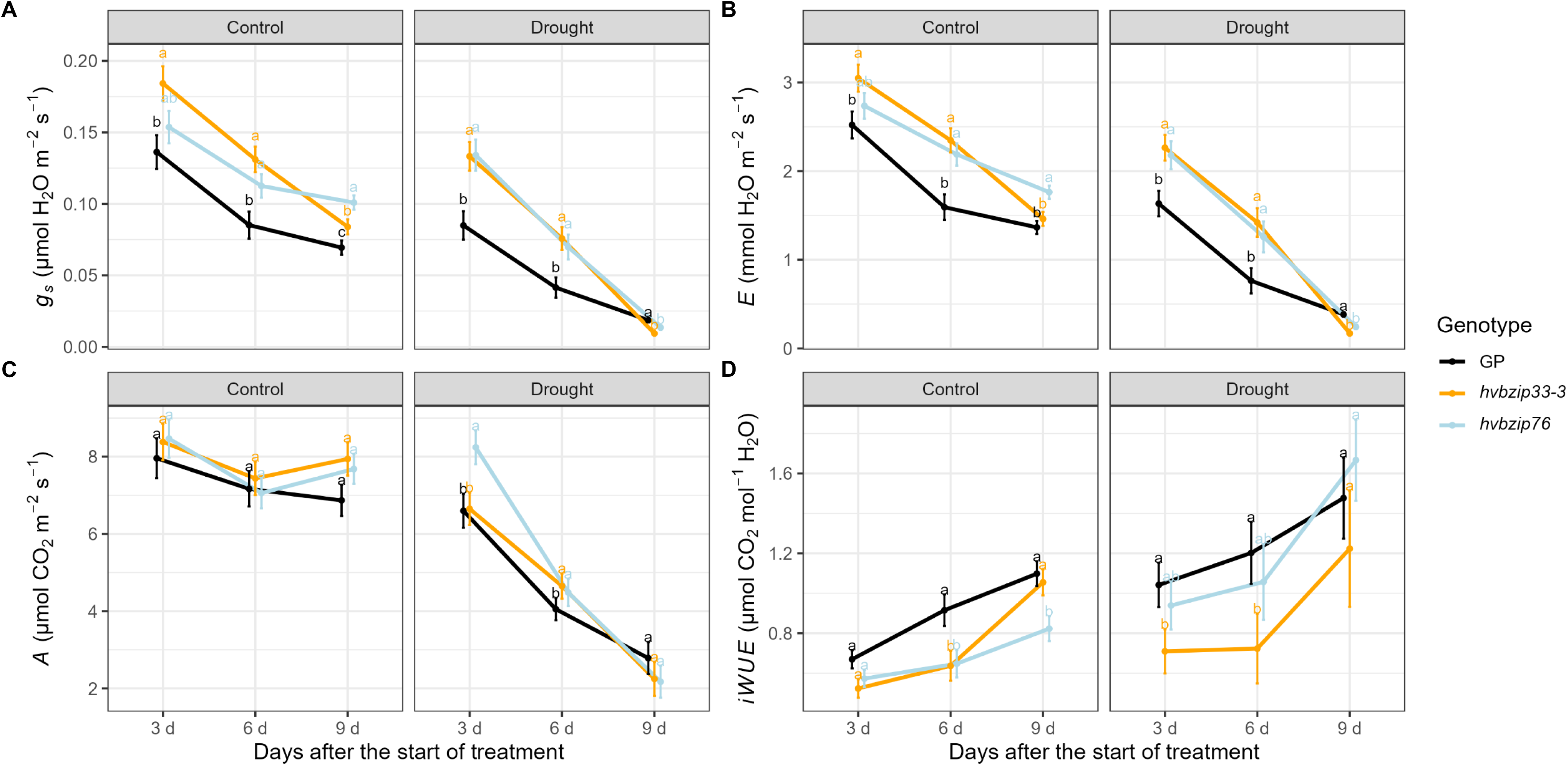
Gas exchange parameters evaluated under control and drought conditions in wild-type Golden Promise (GP) and mutants. The effect of drought stress on (A) stomatal conductance (*g_s_*), (B) transpiration rate (*E*), (C) CO_2_ assimilation (*A*) and (D) intrinsic water use efficiency (*iWUE*). Drought stress was imposed on 14-day-old seedlings via controlled dehydration, ensuring uniform water loss across all experimental units. First gas exchange parameters were evaluated at 3 days after the start of stress treatment (DAS) on the first fully expanded leaf from the top. It was then measured again on 6 and 9 DAS on the same leaf. Two independent experiments were performed. Each experiment comprised four to five biological replications per genotype per treatment. The bar represents mean ± standard error (n = 8-12). Indexed letter represents significant differences (p ≤ 0.05) between the genotypes using the LSD multiple mean comparison test.

### 3.4. Transcriptome analysis of *hvbzip33* and *hvbzip76* mutants in barley

Next, we performed a genome-wide transcriptomic analysis of wild-type GP, *hvbzip76* and *hvbzip33-3* to identify differentially regulated genetic components under control and drought conditions. In accordance with our previous study (Shrestha *et al*., 2022), we observed that *HvbZIP76* expression was upregulated in barley in response to drought stress. We did not detect *HvbZIP33* expression in leaf tissue at the seedling stage under either control or stress conditions (Figure S7).

A total of 19004 genes were co-expressed across all examined inbreds under both control and drought conditions (Figure S8). Principal component (PC) analysis of these global expression patterns revealed that PC1 (46.13% variance) and PC2 (37.18% variance) together accounted for 83.31% of the total transcriptomic variation (Figure 5A). The two mutants clustered together along PC2 and were primarily differentiated from wild-type GP by PC1. While PC1 and PC2 were strongly associated with genotype differences in expression, PC3 and PC4 showed stronger correlations with treatment conditions (Figure 5B), indicating that higher-order principal components capture condition-specific transcriptomic responses (Figure 5D).

**Figure 5:**
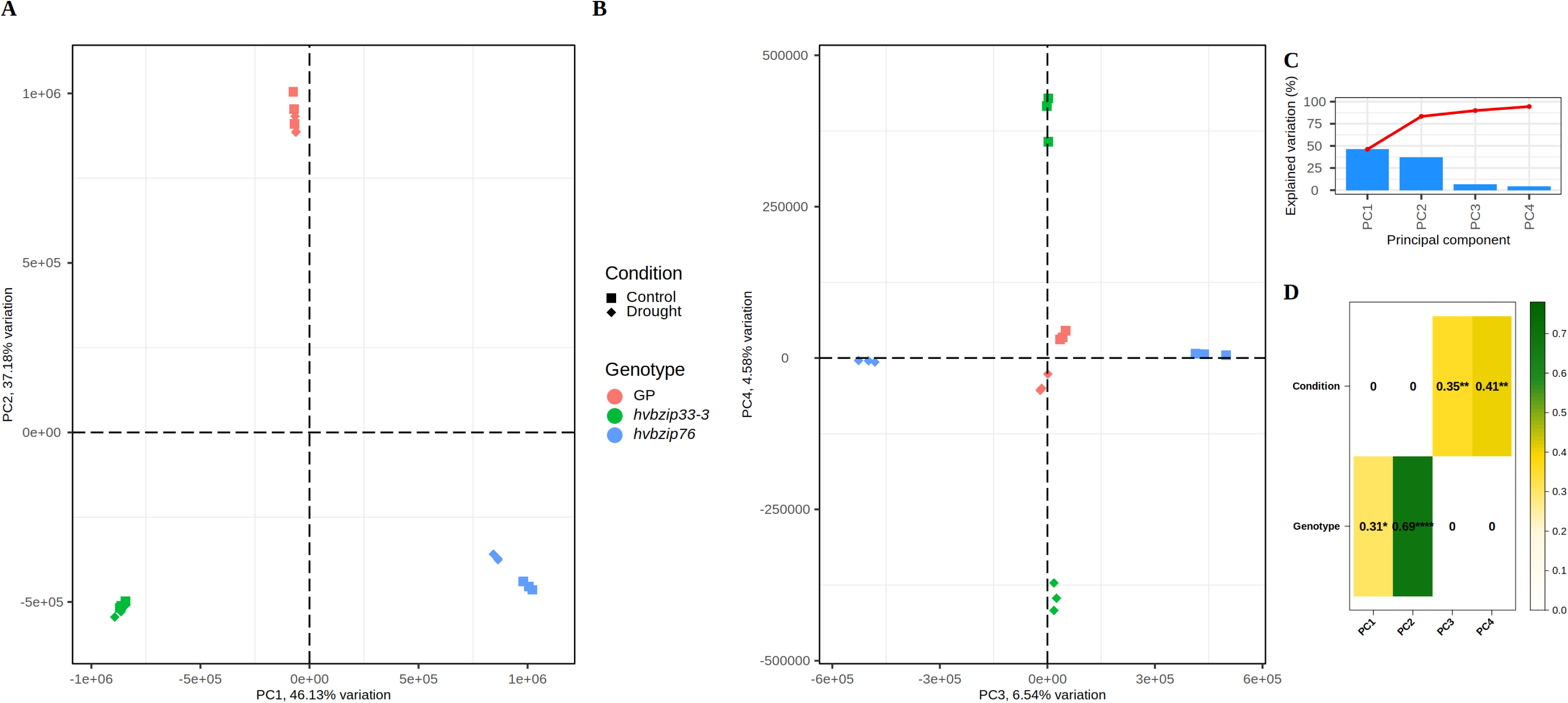
Principal component (PC) analysis of transcriptome profile of GP, *hvbzip33-3* and *hvbzip76*. (A-C) The variance explained by PC1, PC2, PC3, and PC4 on the global gene expression profile of GP, *hvbzip33-3* and *hvbzip76* under control and drought stress. (D) The correlation between PCs and the experimental factors (genotype and growth condition). Asterisks indicate a statistically significant correlation coefficient.

Among the expressed genes, a total of 3368, 3081 and 2448 DEGs were identified between control and drought conditions in *hvbzip76*, *hvbzip33-3* and wild-type GP, respectively, suggesting a higher number of DEGs in mutants than wild-type GP (Figure 6A, Table S3-S10). We detected a substantial central overlap of 1809 genes affected by drought stress, indicating the existence of fundamental drought-stress-responsive pathways. The number of unique DEGs was highest in *hvbzip76* (1230) compared to *hvbzip33-3* (409), indicating (i) broader regulatory influence due to mutation in *HvbZIP76* and (ii) existence of non-redundant regulatory functions. The 777 DEGs between *hvbzip76* and *hvbzip33*-*3* may represent core genes under the control of *HvbZIP76* and *HvbZIP33* (Figure 6A).

**Figure 6:**
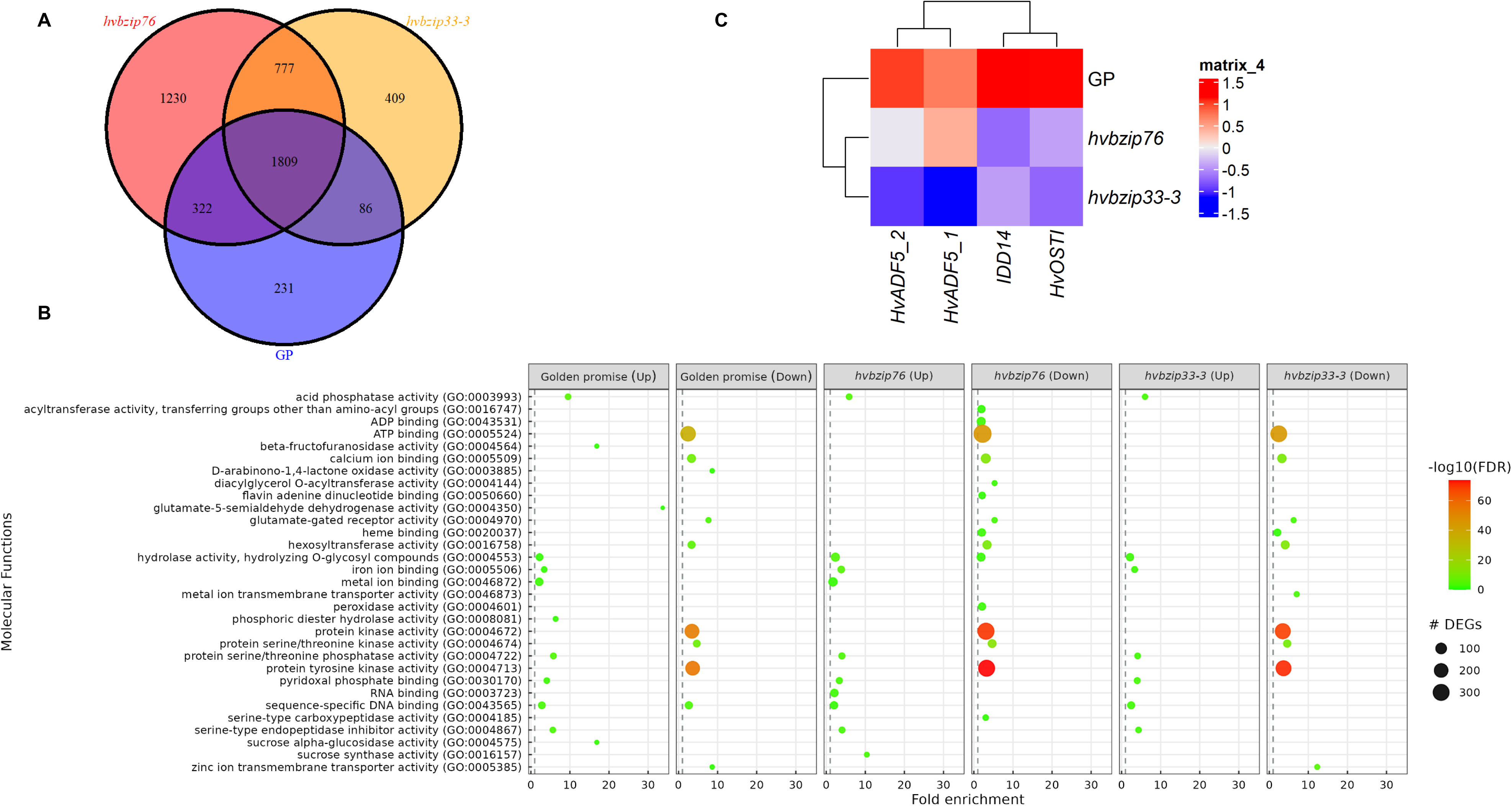
Differentially expressed genes (DEGs) for control versus drought comparison among GP and mutants. (A) Venn diagram illustrating the number of DEGs representing core drought-induced response genes, possible overlapping targets and unique to *HvbZIP33* and *HvbZIP76*. (B) Gene ontology classification and enrichment analysis. The circle’s size indicates the number of genes. The color indicates the *-log10* transformation of the false-discovery-rate p-value. (C) Heat map of fold change (control vs drought) of barley orthologs of genes known to be associated with stomata movement in Arabidopsis and rice. Red and blue color indicate the high and low fold change, respectively under drought compared to control conditions.

GOEA identified 19 biological processes and 38 molecular functions across six differential gene-expression contexts (Figure 6B and Figure S9). The context represents upregulated and downregulated genes in wild-type GP and mutants. The canonical drought-stress response GO terms, such as response to water (GO:0009415), proline biosynthetic processes (GO:0006561), and protein phosphorylation (GO:0006468), were the most enriched among DEGs across all three genotypes. While two of the GO terms mentioned above were associated with genes upregulated under drought stress (GO:0009415 and GO:0006561), we observed coordinated suppression of genes encoding broad protein kinase activity. Interestingly, a specific group of protein kinases (GO:0004672) was linked among the upregulated genes under drought stress. Protein dephosphorylation (GO:0006470) and embryo development (GO:0009790) processes were also associated with up-regulated genes across all the genotypes. One of the most striking differences was the response to desiccation (GO:0009269), which was enriched among upregulated genes only in the wild-type GP (Figures 6B and S9).

Around 30 genes with sequence-specific DNA-binding (GO:0043565) properties were present in both up- and downregulated contexts in wild-type GP. In contrast, they were present only in the upregulated contexts in both mutants. Metal ion binding and transport functions were also enriched among the DEGs. While calcium ion binding (GO:0005509) was universally enriched in downregulated contexts, iron ion binding (GO:0005506) was enriched among upregulated genes. It was noteworthy that heme binding (GO:0020037) GO terms were enriched in the downregulated context in both mutants, while metal ion transmembrane transporter activity (GO:0046873, GO:0005385) was particularly enriched in the downregulated context in *hvbzip33-3*. Peroxidase activity (GO:0004601) was enriched among downregulated genes in *hvbzip76* (Figures 6B and S9).

Because we observed higher *g_s_* and *E* values in mutants than in wild-type GP, we examined in detail the expression of genes known to regulate stomatal opening and closing under drought stress. We found that the fold change (control vs. drought) for the gene annotated as *INDETERMINATE DOMAIN PROTEIN 14* (*IDD14*) was lowest in *hvbzip33-3* and *hvbzip76* compared to wild-type GP. Similarly, the fold change relative to the control for the *OPEN STOMATA 1* (*OST1*) gene was also lower in *hvbzip33-3* and *hvbzip76* than in wild-type GP (Figure 6C). The fold change of another gene associated with stomatal movement, *ACTIN-DEPOLYMERIZING FACTOR 5* (*ADF5*), was lowest in *bzip33-3*, followed by *hvbzip76* and highest in wild-type GP (Figure 6C). These genes are known to regulate stomata pore size and higher expression facilitates stomata closure, reducing water loss during drought stress (Mustilli *et al*., 2002; Qian *et al*., 2019; Liu *et al*., 2022). Therefore, the higher fold change in the expression of the above-discussed genes in wild-type GP might explain the earlier reduction in foliar transpiration in GP than in both mutants.

## 4. Discussion

Our study aimed to characterize barley *HvbZIP* genes that are closely related to the Arabidopsis group A bZIP TF family. Phylogenetic analyses showed that 18 *HvbZIP* genes branched within the Arabidopsis group A bZIP TF, which comprises four ABF1-ABF4-like, DPBF3-DPBF4-like, DPBF1-DPBF2-like and bZIP15-like sub-clusters (Figure S1). *HvbZIP33* (referred to as *ABF1* in earlier studies but closely related to *DPBF2*) cotransfected with ABRC:GUS transactivated GUS in a transient expression assay (Schoonheim *et al*., 2007; Schoonheim *et al*., 2009), indicating a possible role in ABA-mediated transcription activation. *HvbZIP33* is the closest Arabidopsis ortholog of DPBF2 and may play a role in silique development. Similarly, *HvbZIP76* (highly similar to ABF1-ABF4) was activated in four different barley accessions in an ABA-dependent manner, indicating a possible role in ABA-signaling (Shrestha *et al*., 2022). Therefore, we selected *HvbZIP33* (DPBF-like clade) and *HvbZIP76* (ABF-like clade) for functional validation in the current study. For both genes, we generated potential loss-of-function mutations using the RNA-guided Cas9 system (Figures 1, S2 and S3).

### 4.1. Function of *HvbZIP33* and *HvbZIP76* genes in plant development

Previous studies have shown that bZIP transcription factors are involved in multiple critical physiological functions, including stress responses, development and metabolism (Jakoby *et al*., 2002). In addition, the DPBF-like clade in Arabidopsis is known to be involved in reproductive development (Lopez-Molina *et al*., 2003; Mendes *et al*., 2013). Since *HvbZIP33* and *HvbZIP76* are also expressed in the inflorescence (Mascher et al., 2017), we initially examined whether the spikes of the corresponding mutants displayed any developmental defects. We observed that the number of grains per ear and thousand-grain weight were not significantly reduced in *hvbzip76* and *hvbzip33* compared to wild-type GP (Figure S5). While *DPBF2* influences silique development in Arabidopsis, functional studies of the *HvbZIP33* and *HvbZIP76* orthologs in rice have primarily focused on stress response rather than reproductive traits, leaving the effects on grain size and number in monocots largely unexplored (Xiang *et al*., 2008; Tang *et al*., 2012; Chuxin *et al*., 2021). Plant height and ear numbers in adult plants of *hvbzip76, hvbzip33* and wild-type GP were not significantly different from each other (Figure S5). In contrast, plant height and shoot fresh weight were markedly lower in *hvbzip33-3* during the seedling stage. This implies that *hvbzip33-3* plants employ compensatory growth mechanisms that allow them to catch up to wild-type by maturity.

### 4.2. *HvbZIP33* and *HvbZIP76* influence foliar transpiration in barley

Drought experiments at the seedling stage revealed a striking physiological difference among the two mutants and the wild-type GP, where the mutants consistently showed higher *g_s_* and *E* at 3 DAS and 6 DAS under drought stress, resulting in lower *iWUE*. This might have contributed to the more rapid wilting and lower RWC in *hvbzip76* and *hvbzip33* than in wild-type GP (Figure 2A). Our observation was consistent with findings in other species. In rice, lines overexpressing *OsbZIP23*, a group A bZIP TF, exhibited a reduced transpiration rate compared with the wild-type (Xiang *et al*., 2008). Similarly, transgenic Arabidopsis overexpressing *CaADBZ1* (a pepper bZIP TF from the same group A clade) showed enhanced ABA sensitivity and stomatal closure (Choi *et al*., 2025). Furthermore, Arabidopsis *ABF* knockout mutants showed higher water loss rates than wild-type in detached leaf assays (Yoshida *et al*., 2010; Seller and Schroeder, 2023) providing cross-species evidence that group A bZIP TFs might function as positive regulators of stomatal closure under drought stress.

Our transcriptome experiment partially explained these aforementioned phenotypic differences. Under drought stress, mutants showed consistently reduced drought stress-induced expression of key regulatory genes compared to wild-type GP. Specifically, *HvADF5* expression showed the lowest fold-change in *hvbzip33-3*, intermediate in *hvbzip76*, and highest in wild-type GP (Figure 6C). The above observation is consistent with *ADF5’s* role in Arabidopsis as a positive regulator of drought-induced stomatal closure, acting downstream of DPBF transcription factors (Qian *et al*., 2019). Similarly, *HvIDD14* and *HvOST1*, both known to enhance ABA sensitivity and mediate stomatal responses in Arabidopsis (Mustilli *et al*., 2002; Liu *et al*., 2022) showed reduced drought induction in both barley mutants (Fig. 6C). The phylogenetic clustering of *HvbZIP33* with the DPBF clade supports its role in this regulatory module.

Collectively, these findings indicate that *HvbZIP33* and *HvbZIP76* might increase drought tolerance by promoting stomatal closure and water conservation through transcriptional regulation of ABA signaling components. The difference in *g_s_* and *E* was less pronounced under control conditions than drought stress, suggesting that the mutations primarily impair stomatal responsiveness rather than baseline development. However, whether the observed phenotypes result from altered stomatal movement, changes in stomatal size and density, or both remains to be determined. Further anatomical and physiological investigation is needed to distinguish these mechanisms, particularly given that stomata account for over 95% of plant water loss (Hedrich and Shabala, 2018).

### 4.3. Transcriptomic redundancy and divergence among wild-type GP, *hvbzip33* **and *hvbzip76* mutants**

On a genome-wide scale, the number of DEGs comparing wild-type GP versus *hvbzip76* and *hvbzip33-3* was 24-40 times lower than the number of DEGs between control and drought stress conditions (Figure 6A). The quantitative disparity reveals a crucial mechanistic insight that DEGs in wild-type GP versus mutants primarily reflect the contribution of a single bZIP gene. In contrast, the stress versus control comparison captures parallel pathways regulated by stress-responsive multi-gene families that extend beyond the genes studied.

GOEA of DEGs between control and drought-stress conditions revealed that canonical drought-stress-responsive biological processes were commonly enriched across three genotypes, indicating that fundamental stress-responsive mechanisms are conserved even in the mutants. Response to water (GO:0009415) was prominently enriched among the upregulated DEGs (Figure 6B, Figure S9). Protein phosphorylation (GO:0006468), particularly SnRKs, was similarly enriched among drought-stress-responsive genes, consistent with the ABA-SnRK2 signaling pathway, wherein ABA-activated SnRK2 kinases (including SnRK2.6/OST1) phosphorylate multiple downstream targets (Geiger *et al*., 2009; Yoshida *et al*., 2010; Yoshida *et al*., 2015). The proline biosynthetic process (GO:0006561) was also enriched among upregulated genes across all three inbreds, confirming that the fundamental metabolic capacity for osmotic adjustment through proline accumulation (Handa *et al*., 1986; Szabados and Savouré, 2010; Du *et al*., 2023) was also preserved in the mutants.

However, striking genotype-specific differences emerged in other GO term enrichments, revealing divergent regulatory capacities. For instance, heme-binding (GO:0020037) and peroxidase activity-related (GO:0004601) were notably enriched among the downregulated genes in both mutants but not in wild-type GP (Figure 6B, Figure S9). These differences are functionally relevant because heme-binding proteins, including peroxidases and catalase, are the critical components of cellular redox homeostasis and antioxidant defense (Foyer and Noctor, 2005; Eid *et al*., 2024). Their downregulation may explain the marginally higher electrolyte leakage (EL) and malondialdehyde (MDA) content (markers of oxidative stress) observed in the mutants under drought stress. Most strikingly, the desiccation response GO term (GO:0009269) was enriched exclusively among the upregulated genes in wild-type GP (Figure 6B, Figure S9). Such transcriptomic difference suggests a direct or indirect role for *HvbZIP33* and *HvbZIP76* in activating genes that protect cellular structures and macromolecules during severe water deficit (Yoshida *et al*., 2015). The absence of this enrichment in the mutants represents a critical functional gap in drought tolerance.

Additional genotype-specific patterns included metal ion-related molecular functions: Ca^2+^ ion binding (GO:0005509) was universally enriched in the downregulated context, while iron ion binding (GO:0005506) showed differential enrichment patterns across genotypes (Figure 6B, Figure S9). The downregulation of Ca^2+^ binding proteins might indicate a mechanism to reduce intracellular Ca^2+^ buffering capacity, facilitating stomatal closure (You *et al*., 2023). Conversely, metal ion transmembrane transporter activity (GO:0046873) was enriched exclusively among downregulated genes in *hvbzip33-3* mutant under drought stress. Finally, the sequence-specific DNA binding process (GO:0043565) showed bidirectional regulation (both up- and downregulated genes) in wild-type GP, whereas this process was observed only in the upregulated context in both mutants. Therefore, the transcriptional landscape in wild-type GP is more balanced, with representation of transcription factors that are both repressed and activated by drought stress. (Figure 6B, Figure S9).

## 5. Conclusion

The fact that knocking out just two bZIP genes of class A family resulted in significant physiological changes suggests limited functional redundancy among these transcription factors in barley. Further research to characterize the complete family of class A bZIP TFs in barley and their individual contributions to stomatal regulation will provide a more comprehensive understanding of this regulatory network. The observation of increased stomatal conductance and transpiration rate in barley knockout mutants, particularly under drought conditions, underscores the critical role of these transcription factors in regulating plant water relations. By functioning as positive regulators of genes involved in stomatal closure, bZIP transcription factors helped wild-type GP to conserve water during drought. The delicate balance between carbon assimilation and water conservation, regulated by bZIP transcription factors, might represent a key aspect of plant adaptation to water-limited environments, with significant implications for agricultural productivity.

## Authors Contribution

AN, JL and AS conceptualized the study. JK, GH and AS created transgenic lines. JB, CS, THN and AS conducted the experiments. MS, VSB, BS and AS analyzed the data. AS wrote the manuscript with the input from BS, JK, MS, VSB and AN. AN and JL supervised the experiment and acquired the funding for the research.

## Acknowledgement

The authors are grateful to Josef Bauer, Jörg Nettekoven and Thomas Gerhardt from the University of Bonn, who organized logistics for greenhouse experiments. We would also like to thank Sabine Sommerfeld (Plant Reproductive Biology, IPK) for excellent technical support. We also appreciate de.NBI for providing the cloud computing platform. The Graduate School GRK2064 was supported by funding from the German Research Foundation and core funding from the Julius Kühn Institute provided the necessary financial support for the research. Authors acknowledge the use of the Grammarly plug-in to prevent grammatical mistakes.

## Data Availability

The raw sequencing reads from the RNA-seq experiment are available under project PRJEB105244 (accession ID: ERA35397371) in the European Nucleotide Archive. The processed, normalized read-count data are provided as supplementary data. All phenotype data are presented in graphs and tables, and the raw data will also be made available upon request to the corresponding authors.

