## Supplementary material for "*HvbZIP33* and *HvbZIP76* have overlapping roles in foliar transpiration in drought-stressed barley": Figure S

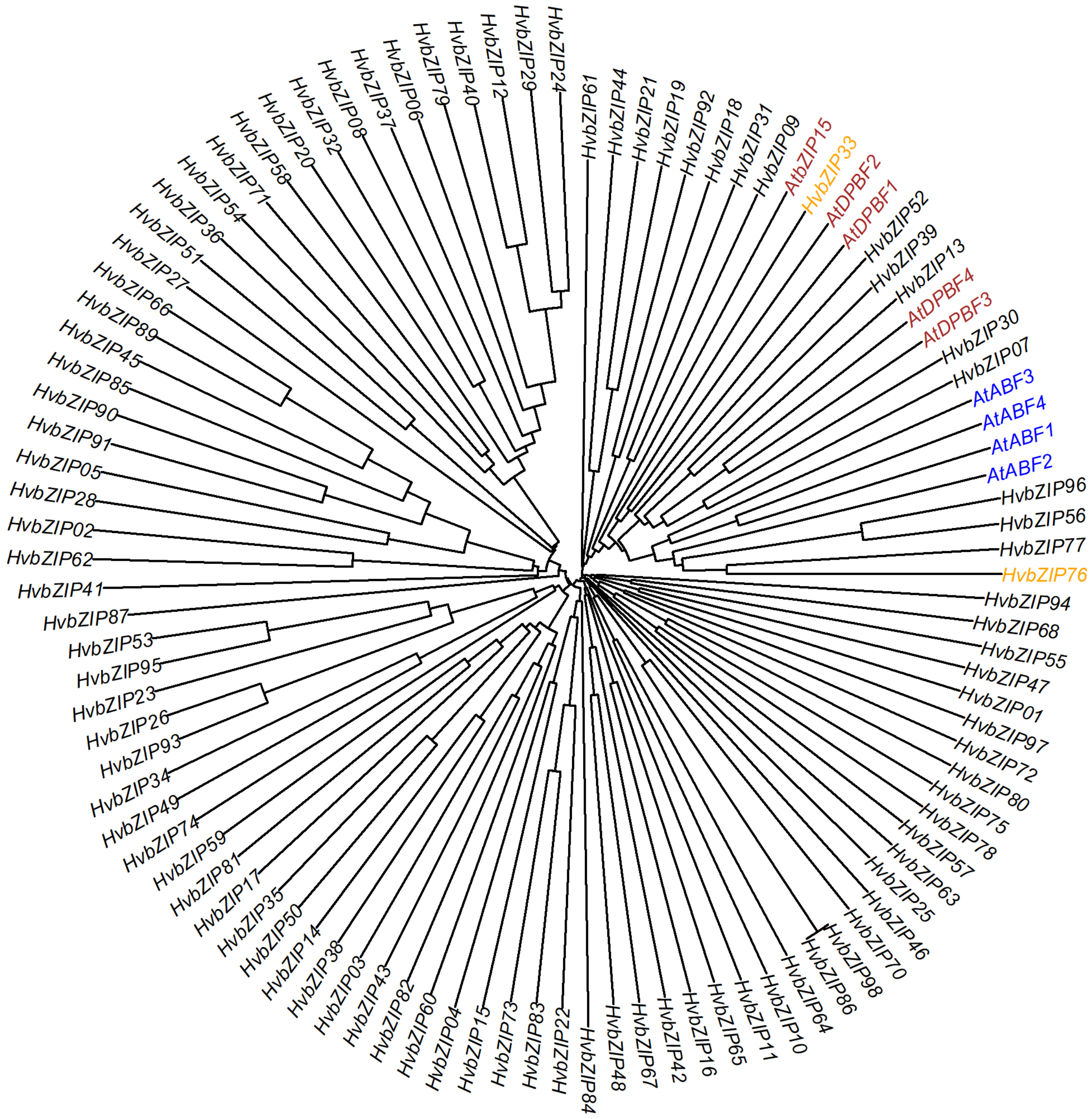


Figure S1: Phylogenetic tree of HvbZIP proteins. Nine class A bZIP transcription factors from Arabidopsis were included in the phylogenetic tree. The genes in blue and brown letters are class A bZIPs identified in Arabidopsis. We have indicated the positions of the two barley bZIP genes, *HvbZIP33* and *HvbZIP76*, in the phylogenetic tree used for functional studies in the current work.

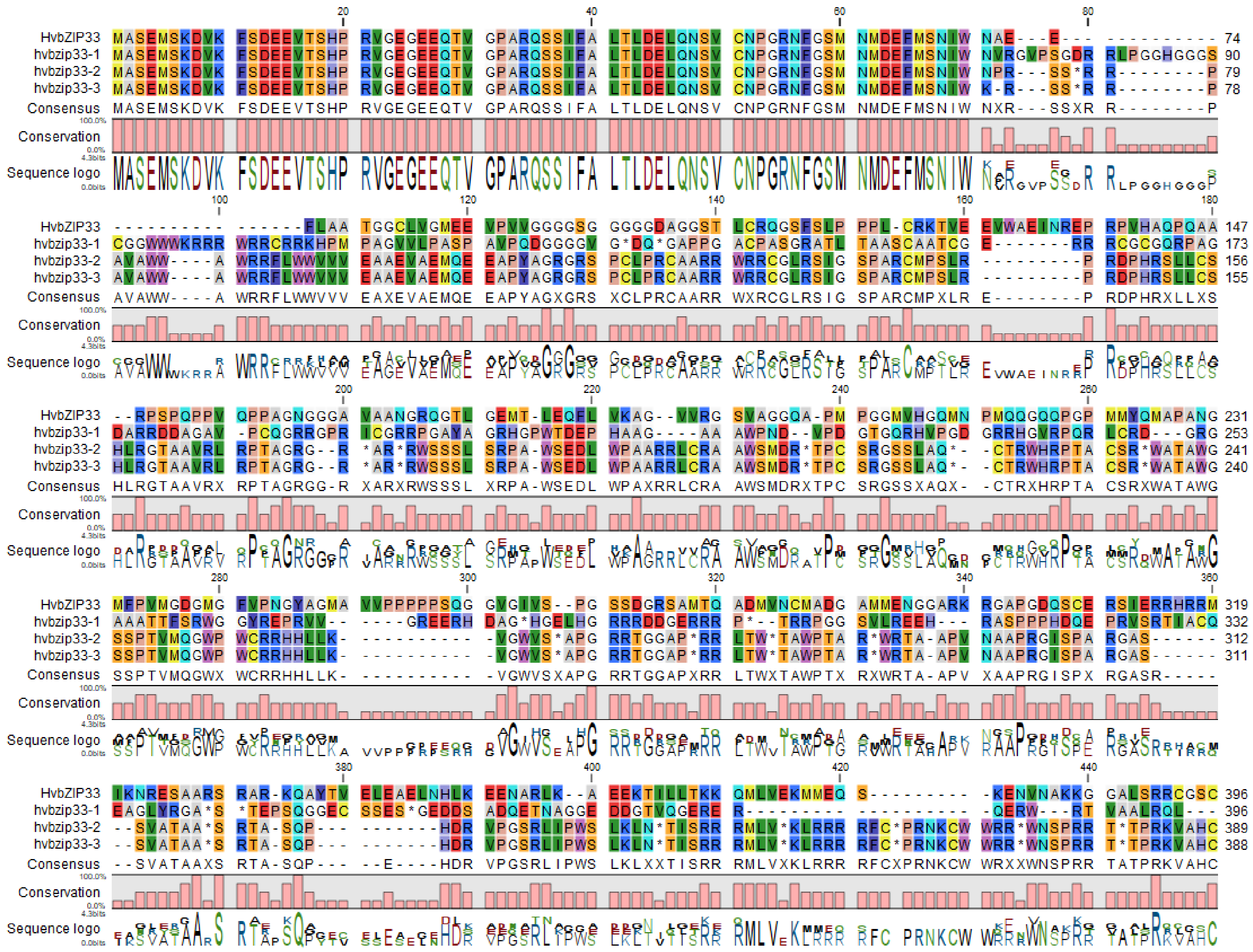

Figure S2: Protein alignment of HvbZIP33 between Golden Promise and mutant lines.

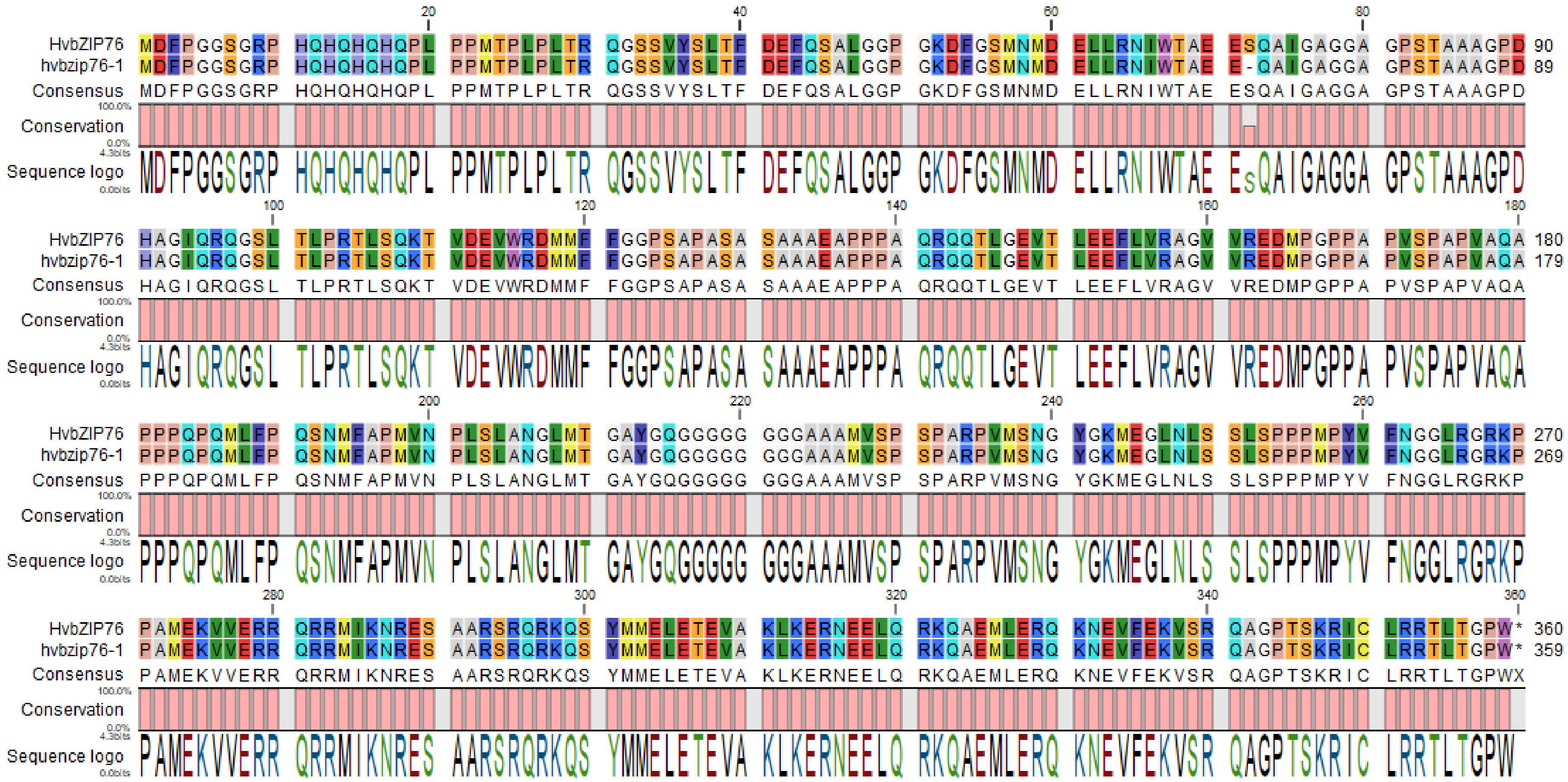

Figure S3: Protein alignment of HvbZIP76 between Golden Promise and the *hvbzip76* mutant line. The conserved domains are amino acid residues 1-32, 77-106, and 275-324.

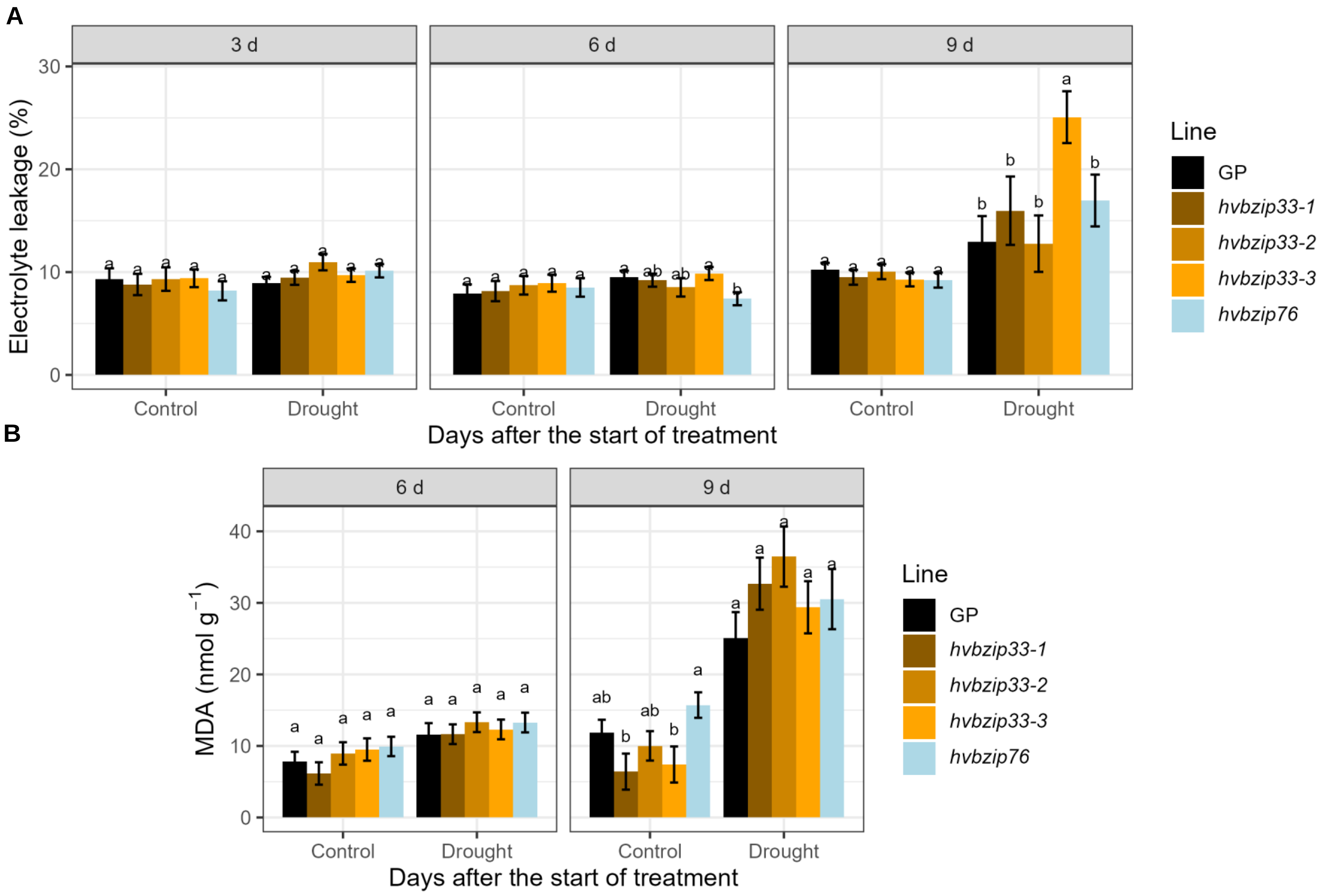


Figure S4: Biochemical traits as stress markers in the leaves of Golden Promise (GP) and mutants. (A) Electrolyte leakage and (B) Malondialdehyde (MDA) content in the first fully expanded leaf from the top at 3, 6 and 9 days after the start of stress treatment (DAS). Drought stress was imposed on 14-day-old seedlings via controlled dehydration, resulting in a uniform water loss across all experimental units. Two independent experiments were performed. Each experiment comprised four biological replicates per genotype and treatment. The bar represents mean ± standard error. Indexed letter represents significant differences ($p \leq 0.05$) between the genotypes using the LSD multiple mean comparison test.

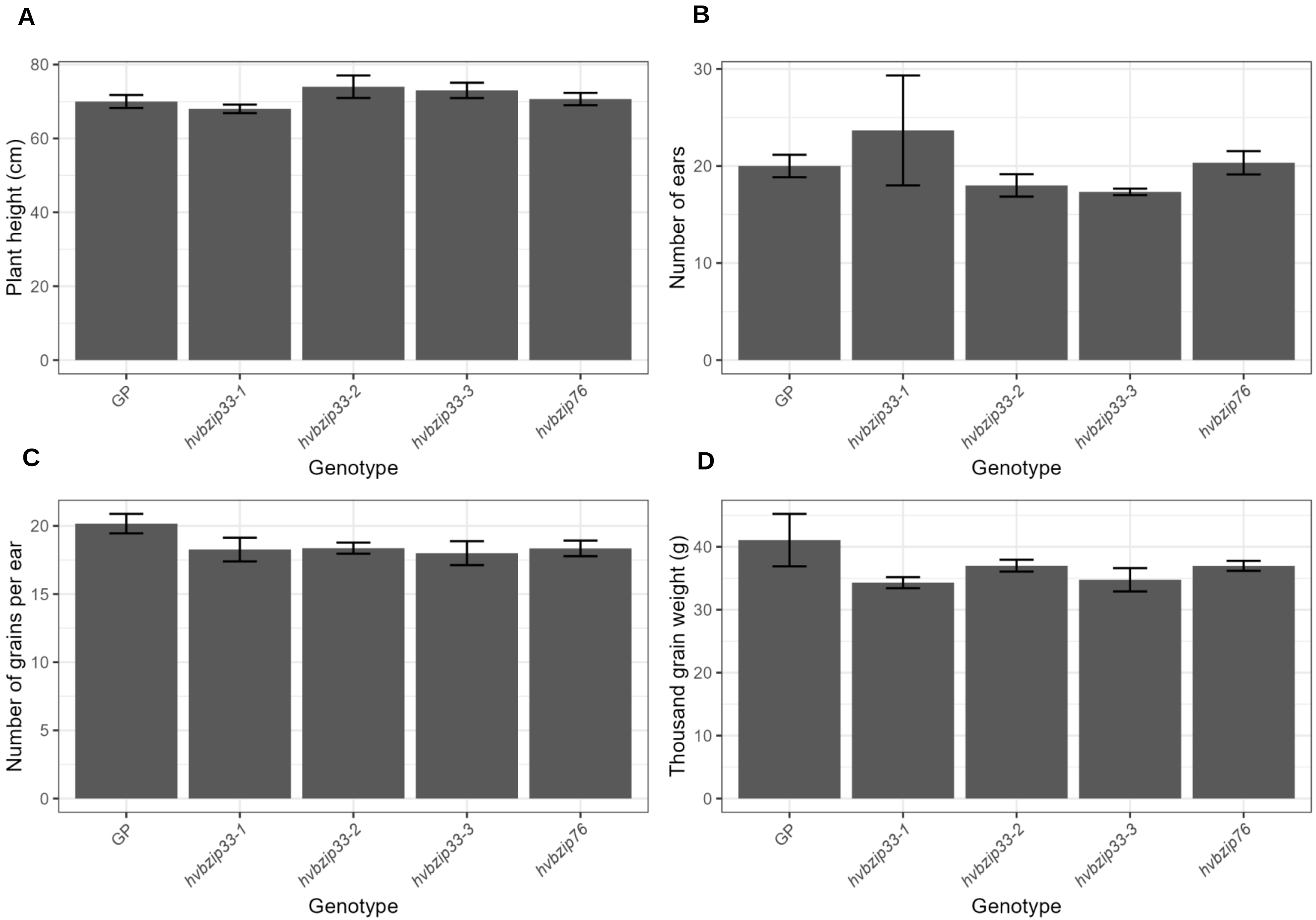


Figure S5: Plant height and spike-related traits in Golden Promise and mutants. Bar represents mean ± standard error (n = 3).

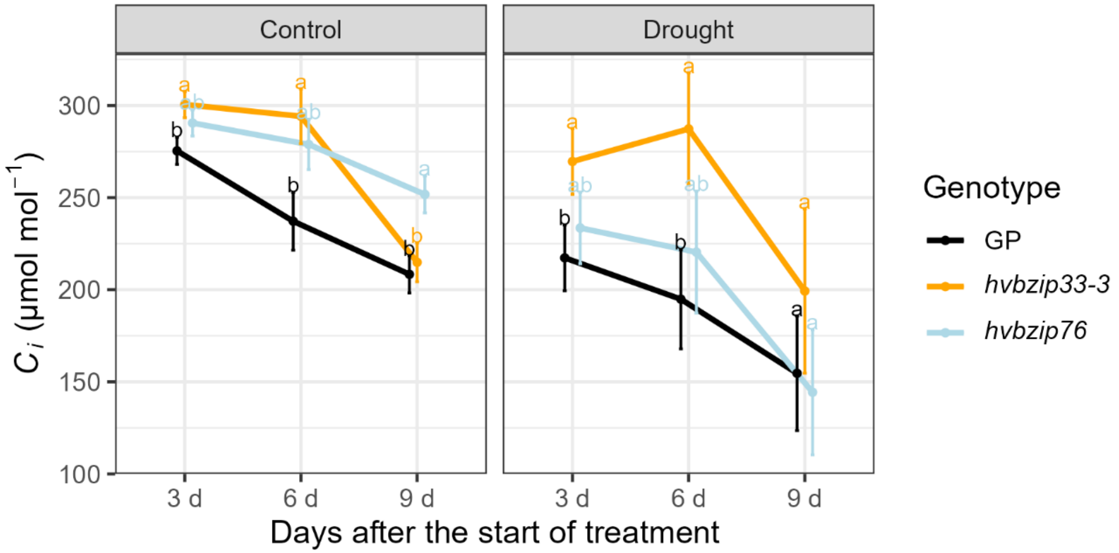


Figure S6: The effect of drought stress on intracellular $CO_2$ concentration ($C_i$) in Golden promise (GP), *hvbzip33-3* and *hvbzip73*. Drought stress was imposed on 14-day-old seedlings via controlled dehydration, resulting in a uniform water loss across all experimental units. First gas exchange parameters were evaluated at 3 days after the start of stress treatment (DAS) on the first fully expanded leaf from the top. Two independent experiments were performed. Each experiment comprised four biological replicates per genotype and treatment. It was then measured again on 6 and 9 DAS on the same leaf. The bar represents mean ± standard error. Indexed letter represents significant differences ($p \leq 0.05$) between the genotypes using the LSD multiple mean comparison test.

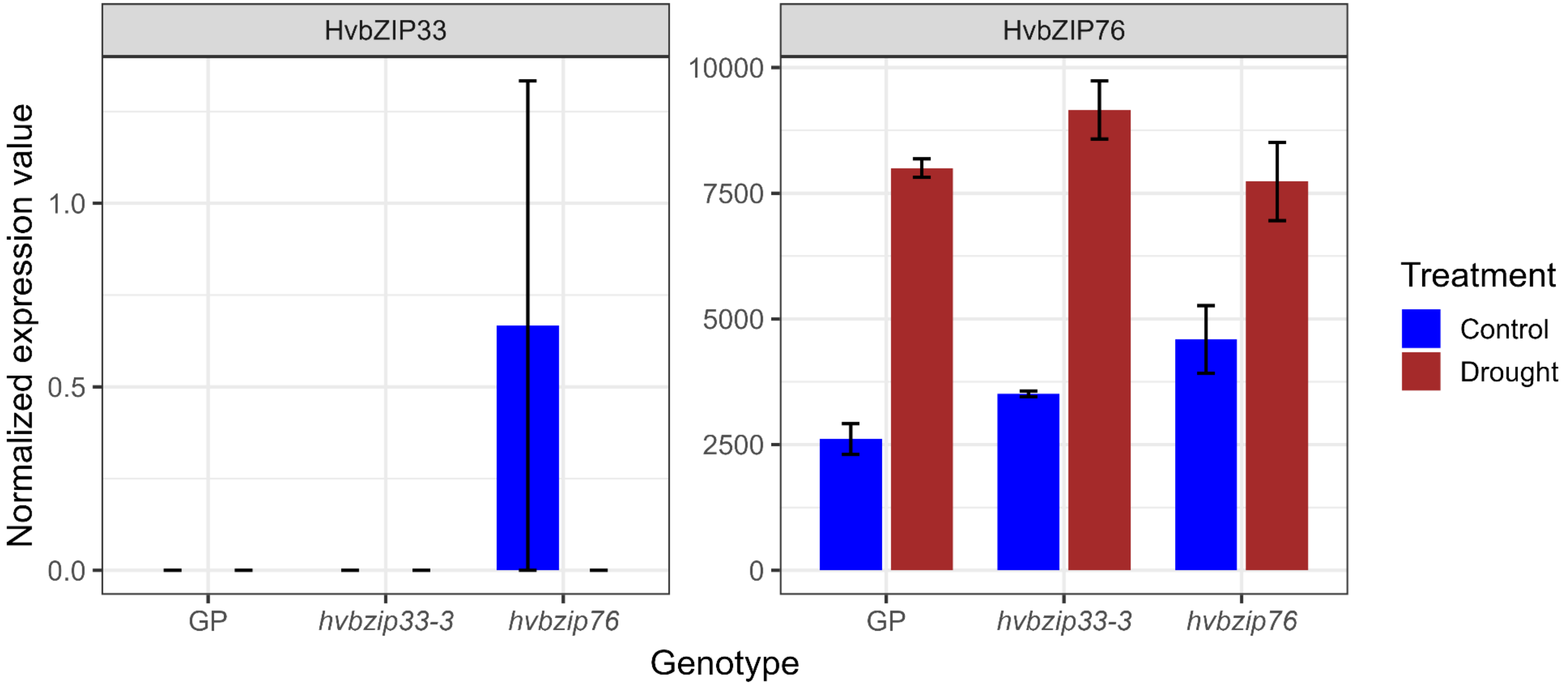


Figure S7: The normalized expression value of *HvbZIP33* and *HvbZIP76* under control and drought conditions.

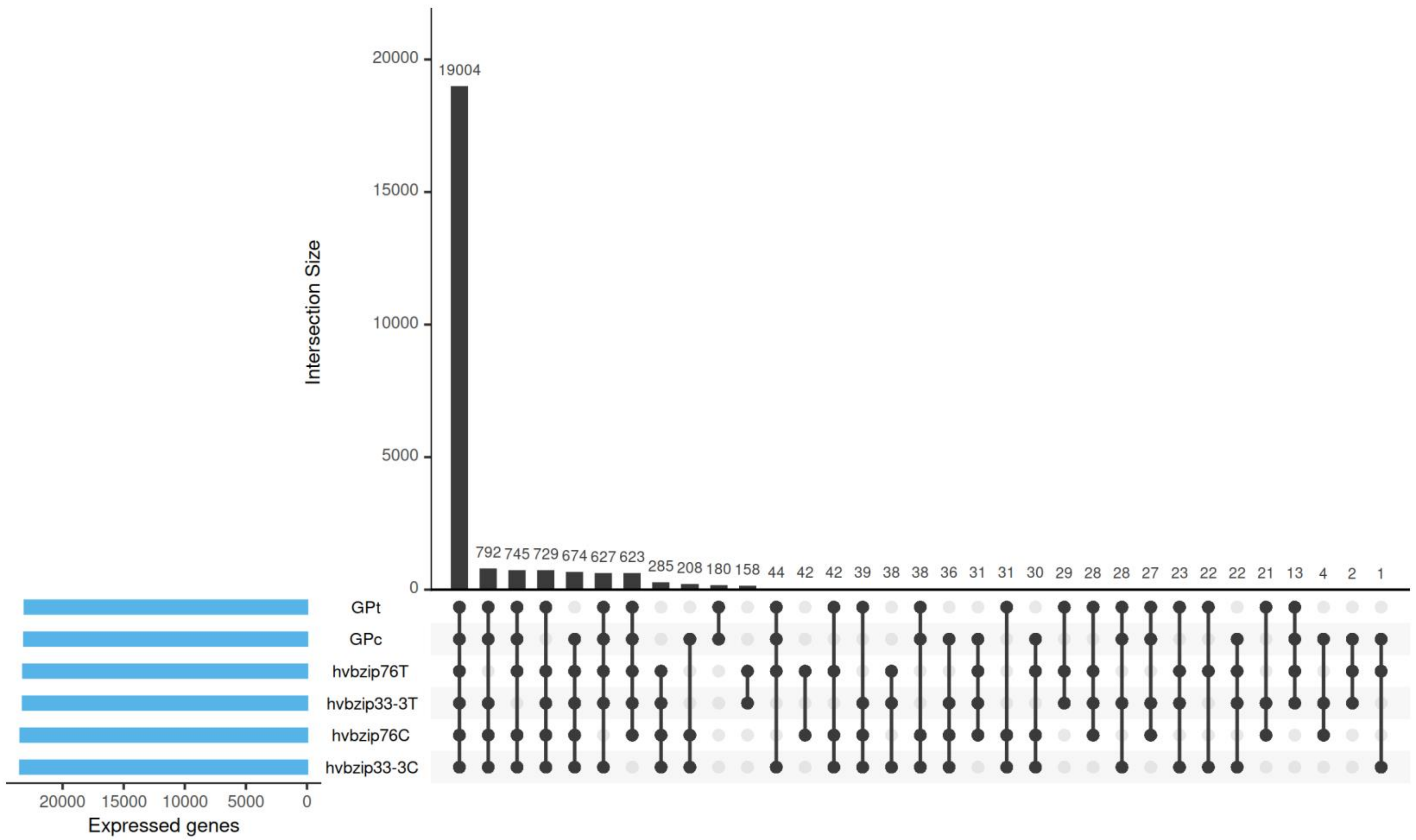


Figure S8: Upset plot of expressed genes across treatment and genetic background. C and T refer to control and drought stress conditions.

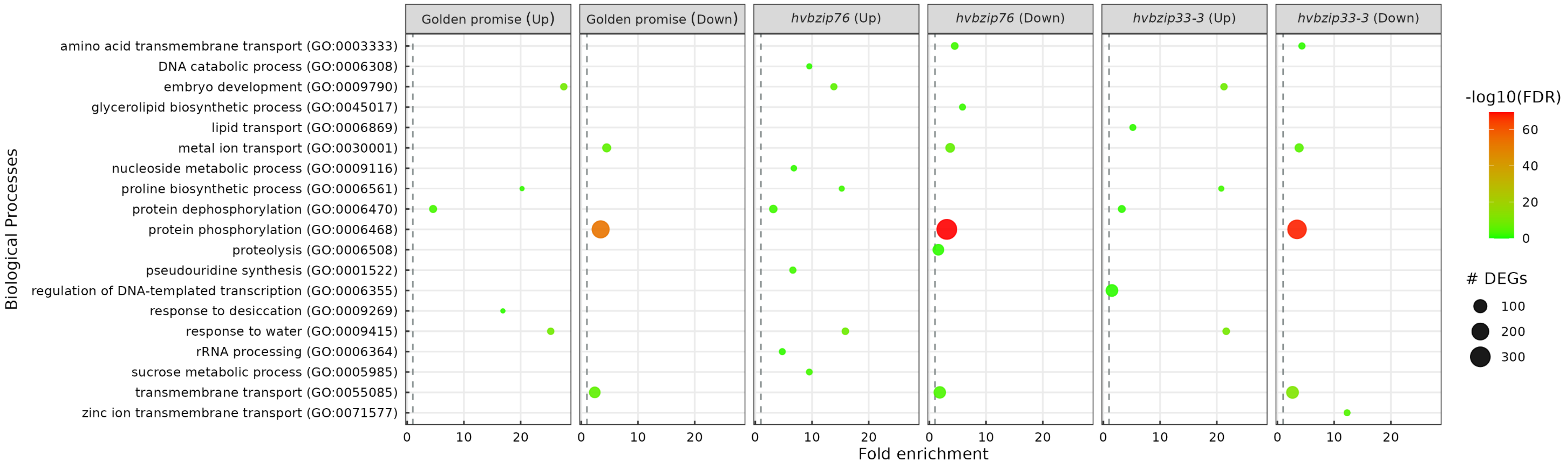


Figure S9: Enriched biological processes based on gene ontology classification of differentially expressed genes between control and drought conditions.
